# Studying early embryogenesis in the flatworm *Maritigrella crozieri* indicates a unique modification of the spiral cleavage program in polyclad flatworms

**DOI:** 10.1101/610733

**Authors:** Johannes Girstmair, Maximilian J. Telford

**Affiliations:** Centre for Life’s Origins and Evolution, Department of Genetics, Evolution and Environment, University College London, London, WC1E 6BT United Kingdom; Max Planck Institute of Molecular Cell Biology and Genetics, Pfotenhauerstraße 108, 01307 Dresden, Germany

**Keywords:** Blebbing, Evo-devo, Light-sheet microscopy, Live imaging, Polyclad flatworms, SPIM, Spiralians, Symmetry breaking, Turbellarians

## Abstract

**Background:** Spiral cleavage is a conserved early developmental mode found in several phyla of Lophotrochozoans with highly diverse adult body plans. While the cleavage pattern has clearly been broadly conserved, it has also undergone many modifications in various taxa. The precise mechanisms of how different adaptations have altered the ancestral spiral cleavage pattern is an important ongoing evolutionary question and adequately answering this question requires obtaining a broad developmental knowledge of different spirally cleaving taxa.

In flatworms (Platyhelminthes), the spiral cleavage program has been lost or severely modified in most taxa. Polyclad flatworms, however, have retained the pattern up to the 32-cell stage. Here we study early embryogenesis of the cotylean polyclad flatworm *Maritigrella crozieri* to investigate how closely this species follows the canonical spiral cleavage pattern and to discover any potential deviations from it.

**Results:** Using live imaging recordings and 3D reconstructions of embryos, we give a detailed picture of the events that occur during spiral cleavage in *M. crozieri*. We suggest, contrary to previous observations, that the 4-cell stage is a product of unequal cleavages. We show that that the formation of third and fourth micromere quartets are accompanied by strong blebbing events; blebbing also accompanies the formation of micromere 4d. We find an important deviation from the canonical pattern of cleavages with clear evidence that micromere 4d follows an atypical cleavage pattern, so far exclusively found in polyclad flatworms.

**Conclusions:** Our findings highlight that early development in *M. crozieri* deviates in several important aspects from the canonical spiral cleavage pattern. We suggest that some of our observations extend to polyclad flatworms in general as they have been described in both suborders of the Polycladida, the Cotylea and Acotylea.

## Background

The Lophotrochozoa is one of two major clades of protostomes, sister group of the Ecdysozoa, (Aguinaldo et al., 1997; Dunn et al., 2008; Halanych et al., 1995; Hejnol et al., 2009; Pick et al., 2010). It contains approximately a dozen morphologically diverse and mostly marine phyla. While the adult morphology of the different phyla gives few obvious clues as to their close relationships, it has long been recognised that a subset of lophotrochozoan phyla share striking similarities in the earliest events of their embryology, most notably in the spatial arrangement of early blastomere divisions, a developmental mode known as spiral cleavage (Hejnol, 2010; Henry, 2014; Lambert, 2010). Representative lophotrochozoan phyla with spiral cleavage comprise annelids, molluscs, nemerteans, flatworms, phoronids and entoprocts (Henry, 2014; Lambert, 2010) and recent phylogenetic results show that these spirally cleaving phyla form a clade within the Lophotrochozoa (Marlétaz et al., 2019). The monophyly of the spirally cleaving phyla strongly suggests a single origin of the spiral cleavage mode. The fact that spiral cleavage has been maintained in these animals since they diverged in the early Cambrian, over half a billion years ago, argues that some selection pressure for maintaining spiral cleavage exists.

There are several aspects of spiral cleavage that appear to be highly conserved. The first is the spiral pattern itself: Embryos of the eight-cell stage consist of four larger vegetal macromeres, 1Q, and four smaller animally positioned micromeres, 1q, each sitting skewed to one side of their sister macromere, above the macromeres’ cleavage furrows. The typical spiral deformations (SD) of macromeres show a helical twist towards one side with respect to the animal-vegetal axis. This is best seen if the embryo is viewed from the animal pole. The resulting spiral shape taken by all four macromeres is either clockwise (dexiotropic) or counter clockwise (laeotropic). In subsequent rounds of division, the larger macromeres again divide unequally and asymmetrically sequentially forming the second and then the third quartets of micromeres. During these divisions the spiral deformations appear in alternating dexiotropic/laeotropic directions (the rule of alternation) up to the fifth cleavage where a 32 cell-stage is reached. At this stage, eight cells of each quarter of the embryo can be traced back to one of the large cells at the four-cell stage and constitute the four quadrants, A, B, C and D. This stereotypical production of quartets means that individual blastomeres can be reliably recognised (and arguably therefore homologised) across spiralian phyla through development. To a variable extent, these homologous blastomeres have been shown subsequently to form lineages with similar fates across the Lophotrochozoa (Henry and Martindale, 1998; Henry and Martindale, 1999; Lyons and Henry, 2014)

The, four quadrants A, B, C and D, that can be recognized in spirally cleaving embryos, and sometimes individually identifiable as early as the four-cell stage, typically correspond to specific body axes. The D quadrant of spiralian embryos has received particular attention from comparative embryologists - once specified, it is involved in major events of embryonic organization. One D quadrant micromere (typically 4d i.e. the micromere descendant of the 4^th^ macromere division) initially and uniquely undergoes a bilaterally symmetrical division determining the dorso-ventral and left right axes of the embryo and then going on to produce large amounts of the future dorsal-posterior part of the embryo. 4d descendants go on to produce endomesoderm (endoderm and mesoderm) (Dorresteijn et al., 1987; Lambert, 2008; Van den Biggelaar and Guerrier, 1983; Verdonk and Van den Biggelaar, 1983). In snails it has been shown that descendants of the D quadrant also possess organizer-like functions (Clement, 1962; Lambert and Nagy, 2001; Martindale, 1986; van den Biggelaar, 1977) The D quadrant lineage arguably holds some of the most conserved features found in spiral cleavers so far.

While spiral cleavage is generally recognized as homologous and highly conserved across spiralian lophotrochozoans, there are, nevertheless, reports of variations on this conserved theme and even complete loss of this mode of development in different species. Alterations to the spiral cleavage mode include unusual arrangements and differences in relative sizes of blastomeres, alternative cell fates including rare derivations of the otherwise highly conserved origin of the mesoderm (Meyer et al., 2010), and even complete loss of the spiral arrangements of blastomeres (Hejnol, 2010).

The mesoderm arises from the D quadrant and, although its lineage is conserved in lophotrochozan development, there are two different ways of specifying which of the four quadrants is the D quadrant. This crucial step can either be achieved early in development by producing blastomeres of different sizes (and presumably containing different maternal transcripts or proteins). Such embryos are classified as “unequal cleavers” whereby the D blastomere at the 4-cell stage is typically the largest cell (Freeman and Lundelius, 1992; Lambert and Nagy, 2003). In other species (equal cleavers), D-quadrant specification is thought to take place by an inductive interaction, usually between one of the large macromeres and the first quartet of micromeres (see Lyons and Henry, 2014). In the latter case, the specification of the D quadrant occurs later in development (Freeman and Lundelius, 1992), with some significant variations in timing.

To reconstruct the ancestral features of spiral cleavage and to further the understanding of the adaptive basis of any modifications of the spiral cleavage program, it is essential to broaden our knowledge of the phylogenetically conserved and variable features of the spiral cleavage program by studying the full diversity of spiral cleavers. Here we focus on both the conserved and the derived aspects of early spiral cleavage in one important but understudied lophotrochozoan phylum: the Platyhelminthes (flatworms). Across the Platyhelminthes, a wide range of different evolutionary developmental modes is found, indeed, in most members of the phylum spiral cleavage has been lost entirely. Only the Polycladida and its sister group, the Prorhynchida (Egger et al., 2015; Martín-Durán and Egger, 2012), have retained an apparently canonical form of spiral quartet cleavage. For this reason, both taxa are excellent candidates for evolutionary comparative studies (Lapraz et al., 2013; Martín-Durán and Egger, 2012).

Most of our current knowledge of polyclad embryogenesis derives from observations made in embryos of *Hoploplana inquilina* (Boyer et al., 1998; Surface, 1907), which belongs to the Acotylea, one of the two major suborders found within polyclad flatworms. Here, we investigate the early cleavages of *Maritigrella crozieri* a member of the Cotylea - the second major clade of polyclads. *Maritigrella* has been recently introduced as a model to study flatworm evolution and development (Girstmair et al., 2016; Lapraz et al., 2013; Rawlinson, 2010). We provide the most detailed description to date of the early development of a cotylean polyclad flatworm. To visualize the development of embryos *in vivo* we used a recently established live-imaging setup, using selective plane illumination microscopy (SPIM) via the OpenSPIM open access platform (Gualda et al., 2013; Pitrone et al., 2013), which allows *in vivo* recordings and precise 3D reconstructions of polyclad flatworm embryos (Girstmair et al., 2016). We use 4D live imaging to visualize details of the early development of *M. crozieri* and we examine cell volume measurements of blastomeres from the first and second cleavages. Live imaging also allows us to make new observations of blastomere dynamics during spiral quartet cleavage.

## Results and Discussion

### Overview of the spiral cleavage pattern in *Maritigrella crozieri*

The development of polyclad flatworms closely follows the conserved spiral cleavage mode and this is true of both polyclad suborders, the Acotylea and Cotylea, as well as in direct and indirect developers within both suborders (Boyer et al., 1998; Gammoudi et al., 2011; Goette, 1881; Kato, 1940; Lang, 1884; Lapraz et al., 2013; Malakhov and Trubitsina, 1998; Martín-Durán and Egger, 2012; Rawlinson, 2010; Surface, 1907; Wilson, 1898). Cleavage in polyclads, as in other spiralians, begins with two meridional divisions (from animal pole to vegetal pole) resulting in four cells arranged around the central animal-vegetal axis and these blastomeres have the standard names of A, B, C, D. The stereotypical polyclad cleavage pattern after the four-cell stage from the third to the fifth cleavage (32-cell stage) is summarized in Figure 1, A-C. Thereby three quartets of ectodermal micromeres (1q-3q) are budded at the animal pole by repeated divisions of the large macromeres. In most spiral cleavers a fourth and sometimes even a fifth quartet of blastomeres are formed in this specific geometry. In polyclad flatworms, however, the fourth quartet significantly deviates from the stereotypic cleavage in terms of both relative size of micromeres and macromeres and their orientation. In contrast to the formation of the first three quartets of micromeres, the fourth quartet ‘micromeres’ are considerably larger than the four sister ‘macromeres’ which form as four tiny cells at the vegetal pole (see Figure 1 D). This unusual characteristic of large 4^th^ quartet micromeres has been previously shown in polyclad flatworms including both *H. inquilina* (Boyer et al., 1998) and *M. crozieri* (Rawlinson, 2010).

**Figure 1.**
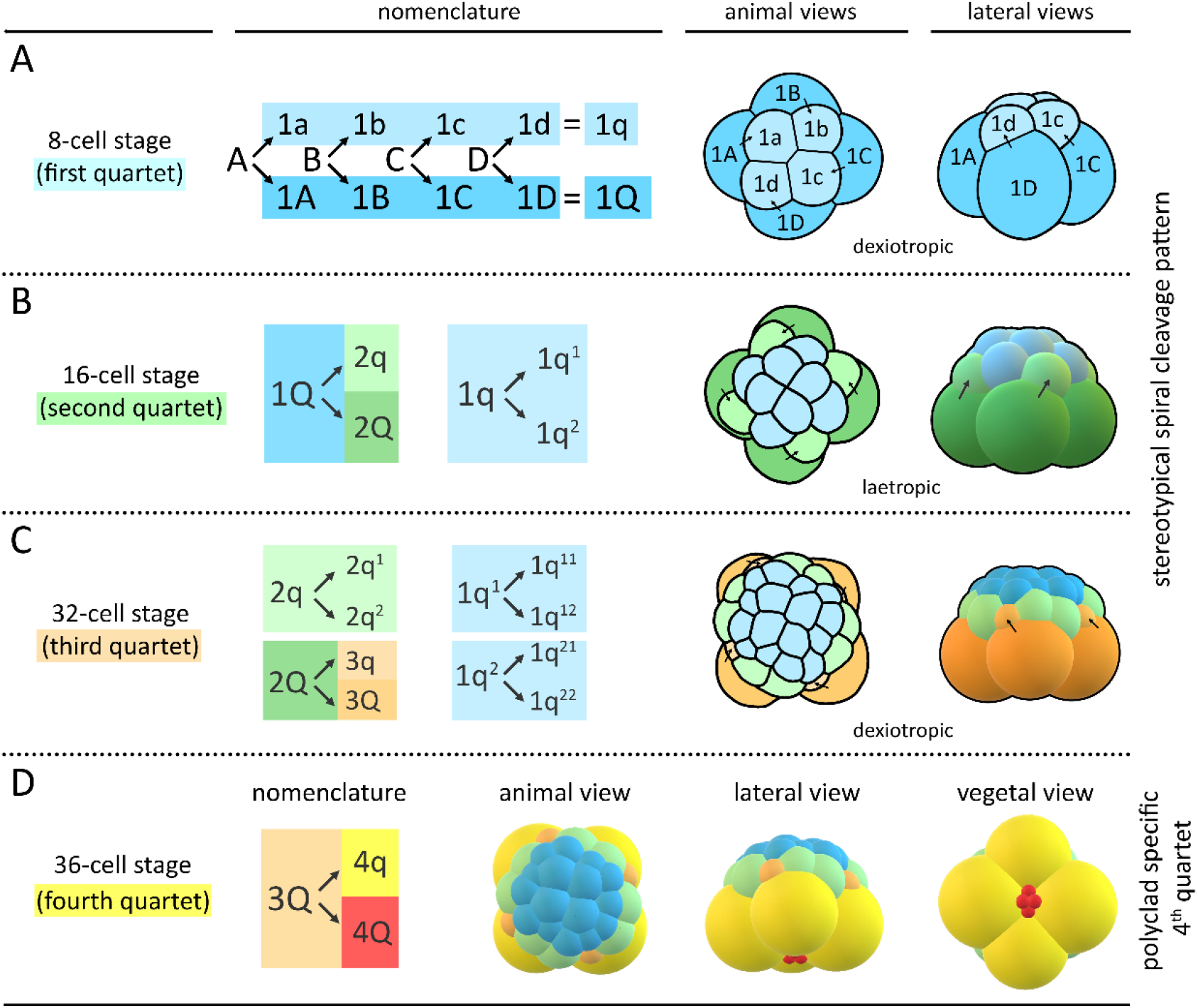
Schematics and nomenclature of the spiral quartet cleavage as found in polyclad flatworms. Micromere and macromere quartets (q and Q, respectively) are colour-coded. **(A)** The third cleavage (4-to 8-cell stage) is unequal and asymmetric. The eight-cell stage embryo consists of four larger vegetal macromeres 1Q, and four smaller animally positioned micromeres 1q sitting skewed to one side of their sister macromere, above the macromeres’ cleavage furrows. The typical spiral deformations (SD) of macromeres show a helical twist towards one side with respect to the animal-vegetal axis. This is best seen if the embryo is viewed from the animal pole. The resulting spiral shape taken by all four macromeres has been shown to be either clockwise (dexiotropic) or counter clockwise (laeotropic) among different lophotrochozoans. In the polyclad *M. crozieri* it is dexiotropic. Notably it has been demonstrated that the mechanism of spiral deformations depends on actin filaments rather than on spindle forming microtubules (Shibazaki et al., 2004). **(B-C)** In subsequent rounds of division, the larger macromeres again divide unequally and asymmetrically sequentially forming the second and then the third quartets of micromeres. During these divisions the spiral deformations appear in alternating dexiotropic/laeotropic directions (the rule of alternation) up to the fifth cleavage where a 32 cell-stage is reached. Up to this point, polyclad flatworms represent a classic example of stereotypic lophotrochozoan spiral quartet cleavage. **(D)** The formation of the fourth cleavage (4Q and 4q) deviates from the typical pattern seen in other spiral-cleaving embryos insofar as the micromeres 4q become large and the macromeres 4Q diminutive. Q = A, B, C, D; q = a, b, c, d.

Our observations of *M. crozieri’s* earliest cleavage pattern, which include live-imaging recordings (Figure 2) and scanning electron microscopy images (Figure 3, A-F) are in accordance with previous 4D recordings up to the 16-cell stage (Lapraz et al., 2013) and descriptions of fixed specimens (Rawlinson, 2010). In some specimens we noted that second cleavages were slightly asynchronous, which explains the occasional observation of embryos in a 3-cell stage before the formation of four similarly sized blastomeres takes place. The characteristic cleavage pattern and spiral deformations are prominent; the 4-to 8-cell transition is dexiotropic (compare Figure 1, A Figure 3, A and Additional file 1). As the division of the first quartet micromeres (1a-1d) is slightly delayed relative to the division of their sister macromeres (1A-1D), an intermediate 12-cell stage forms (Figure 3, C and D). During the generation of new quartets by division of macromeres, the micromeres of the existing quartets also divide and, after the third quartet is completed, the embryo reaches a 32-cell stage.

**Figure 2.**
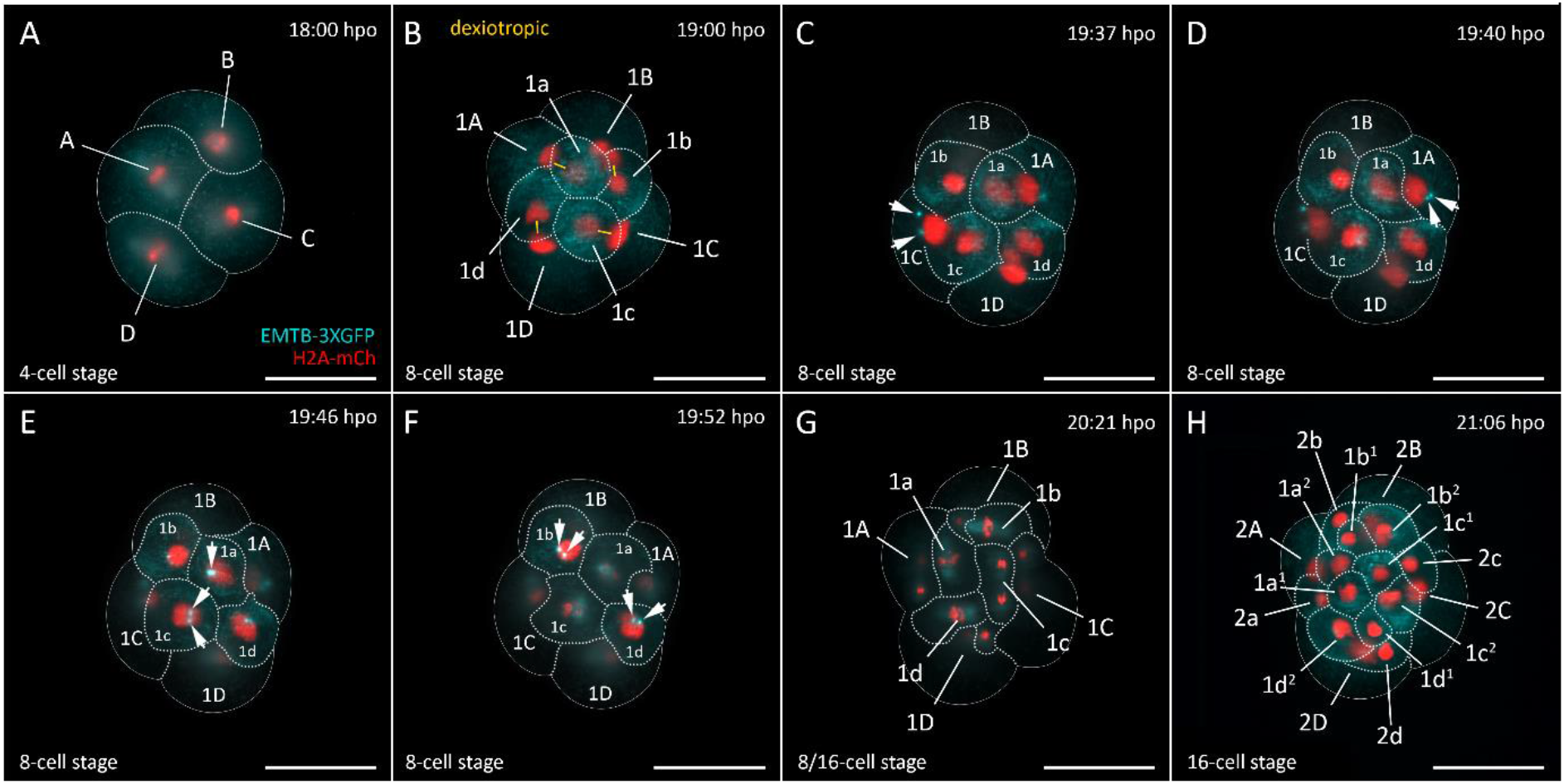
Live-imaging of the transition from a 4-cell stage embryo to a 16-cell stage in *M. crozieri*. **(A)** 4-cell stage with pronounced animal and vegetal cross-furrow cells. (B-F) 8-cell stage preparing for the fourth cleavage round. White arrows point to the appearance of microtubule organizing centre (MTOC) **(G)** Divisions of the fourth cleavage round. (H) The embryo has reached the 16-cell stage is reached; hpo = hours post oviposition. Images captured with an OpenSPIM. Scalebar is 100 μm.

**Figure 3.**
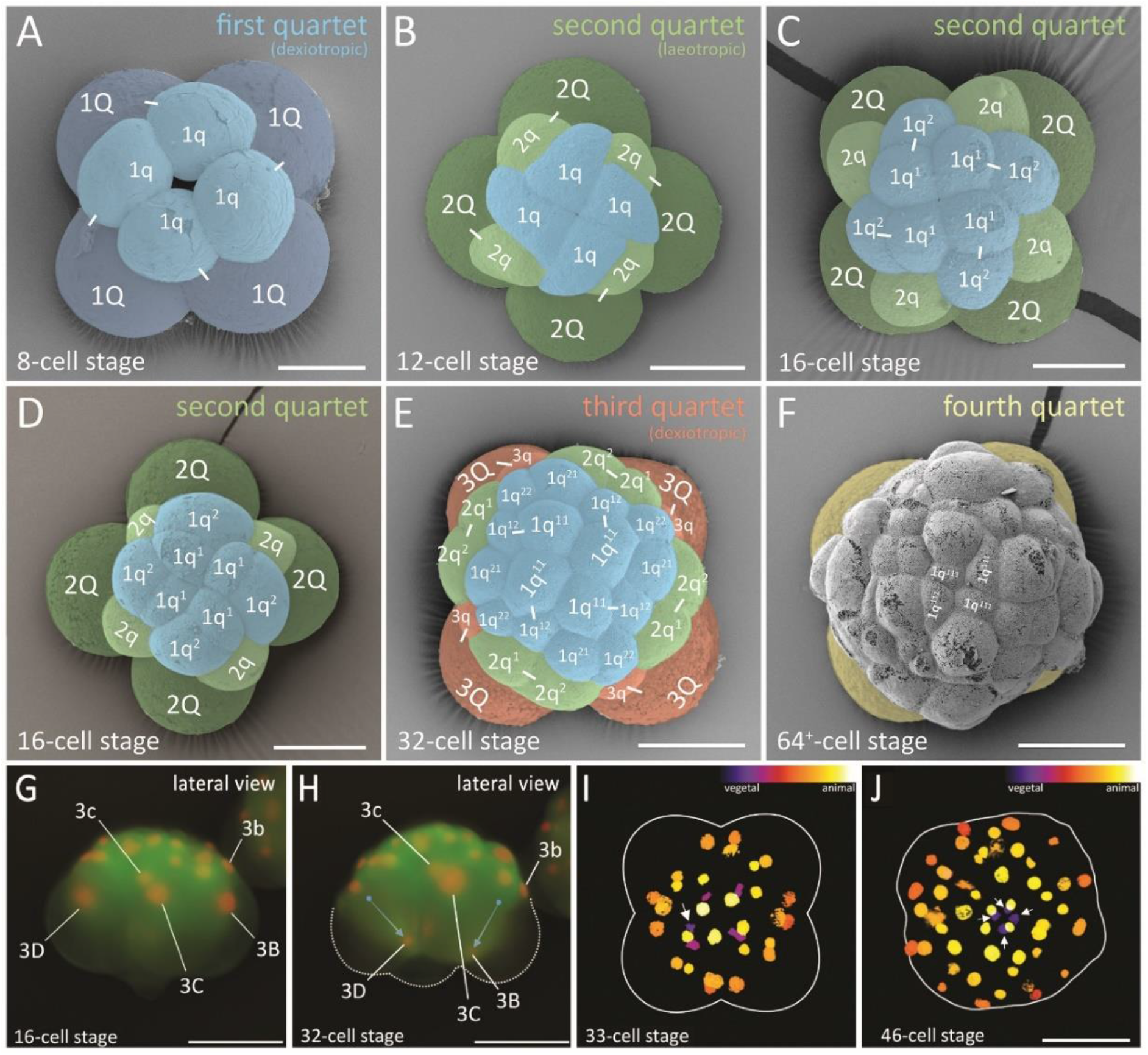
Formation of the four quartets in *M. crozieri*. **(A-D)** SEM pictures coloured according to micromere quartets. **(A)** First quartet (1Q and 1q indicated in blue). **(B-D)** Second quartet (2Q and 2q) indicated in green. **(E)** Third quartet (3Q and 3q) indicated in orange. **(F)** Large fourth micromere quartet (4q) indicated in yellow. G-J: Formation of the fourth quartet **(G)** The 16-cell stage shows macromeres 3B-D and their nuclei at an animal position within the large blastomeres. **(H)** Same embryo as in G but at the 32-cell stage. Nuclei of 3B and 3D are now positioned at the vegetal pole of the macromeres. **(I)** 33-cell stage of a 3D reconstructed embryo (Their depth in the embryo is coded by colours as seen in top right part of the panel. Division of one of the four macromeres (3Q) into 4Q/4q has taken place. The white arrow indicates the newly formed small macromere of the fourth quartet (4Q) coloured purple indicating it is close to the vegetal pole. **(J)** 3D reconstructions showing that all four macromeres comprising the fourth quartet are now positioned at the most vegetal pole of the embryo (coloured purple and indicated by arrows). Scalebar in A-J = 50 μm.

We suggest that, during the polyclad-specific fourth quartet formation, the unusual asymmetric division resulting in large micromeres and small macromeres is achieved in part by a significant displacement of the nuclei in all four macromeres (3Q) prior to their division as is shown here in embryos of *M. crozieri* (Figure 3, G-J). The macromere nuclei, which are typically placed towards the animal pole, shift significantly towards the vegetal pole in 3A-3D (Figure 3, G and H, blue arrows). As a result of these movements, the nuclei of 3A-3D meet at the vegetal pole of the embryo, just before the macromeres divide (Figure 3, I, purple nuclei). The newly formed large micromeres retain most of their size and all the yolk. After the completion of the fourth quartet of micromeres, embryos have reached the 36-cell stage. In polyclads, except for micromere 4d, cells of the fourth quartet do not appear to undergo any further divisions for as long as they can be traced during epibolic gastrulation (Boyer et al., 1998; Rawlinson, 2010; Surface, 1907). At the point when cilia form on the epidermis and embryos start to rotate, cells become difficult to identify and their fates obscure, however, there is evidence from our live imaging recordings that, during epiboly and after bilateral symmetry is established, these small macromeres could engage in further cell-cell interactions. The nuclei of the small macromeres (4A-4D) can be seen in close proximity with nuclei of descendants of micromere 4d^1^ (probably micromere 4d^11^) as is shown in Figure 4 and as a movie (see Additional file 2). This observation suggests that the fourth quartet macromeres undergo later cell interactions and this might play a more important developmental role than has previously been appreciated.

**Figure 4.**
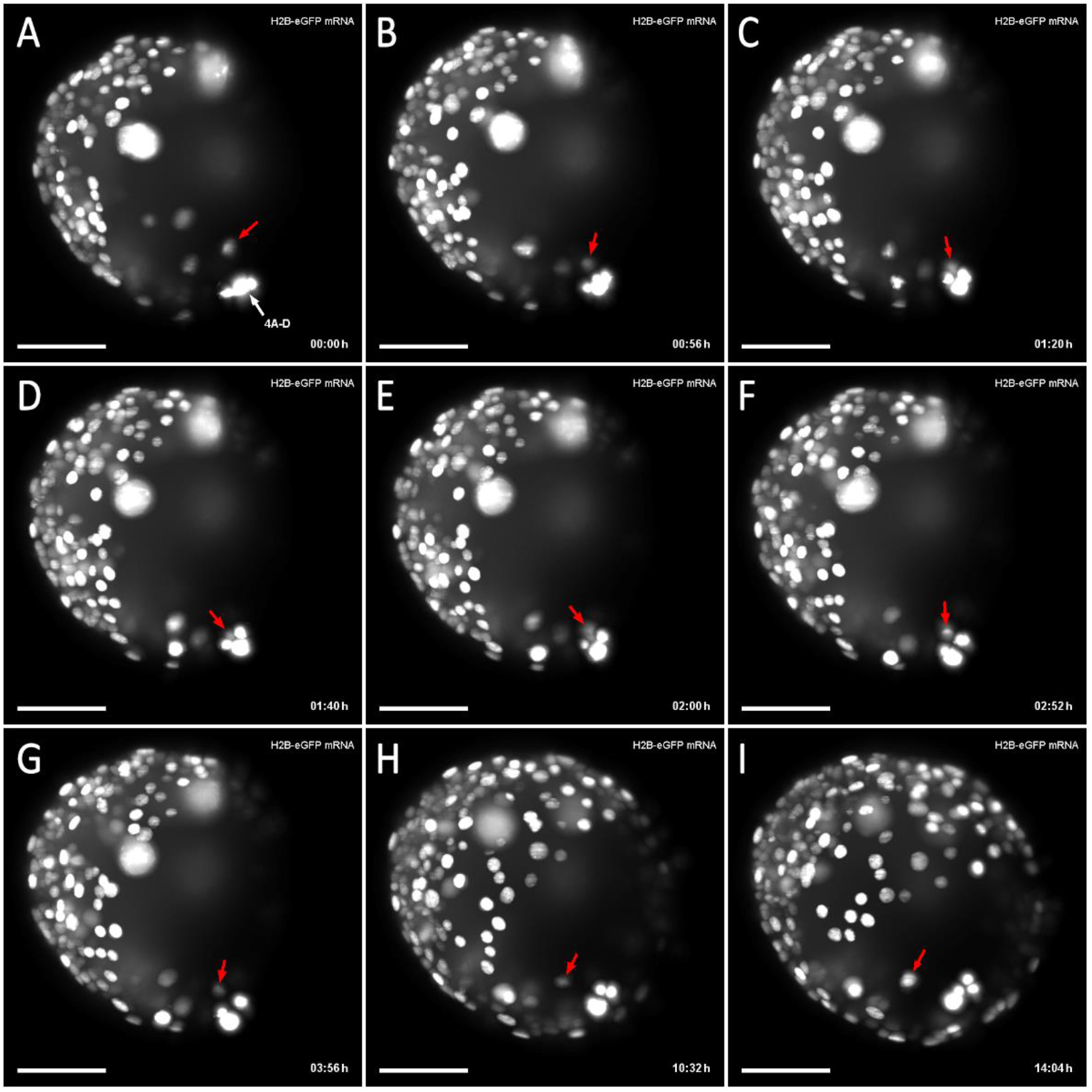
Putative cell-cell interactions observed in the gastrulating polyclad flatworm *M. crozieri*. **(A-I)** A descendant of cell 4d^2^ (red arrow) is traced and can be seen approaching and later departing from small macromeres 4A-D before epiboly is completed. Time represents hours (h) of time-lapse imaging; h is hours of imaging with an OpenSPIM.. Scalebar = 50 μm.

The dramatic changes in cell behaviour from an animally-positioned cleavage position into a vegetal one resulting in small ‘macromeres’ of the fourth quartet are not widely observed in other spirally cleaving embryos. This deviation from typical quartet formation pattern raises the question as to how and why such a modification evolved. Interestingly, in the common bladder snail *Physa fontinalis* the fourth quartet emerges in a very similar way to polyclad flatworms, producing a rosette of four smaller macromeres (4A-4D) at the most vegetal pole and four larger micromeres (4a-4d) above it (Wierzejski, 1905). In *P. fontinalis*, unlike polyclad flatworms, macromere 3D divides earlier in the snail than its sister cells (3A-3C) giving rise to micromere 4d (the mesentoblast). Furthermore, in *P. fontinalis*, cells of the small macromere rosette (4A-4D) undergo a further division producing a fifth quartet of micromeres through equal divisions of 4A-4C.

### The four-cell stage is a product of asymmetric cleavages in *M. crozieri*

In spiral cleavers, equal and unequal cleavage types can be readily distinguished during the first two divisions. The cleavage mode has been thought to reflect the way in which the embryo determines one of its four quadrants to become designated as the D quadrant. (Arnolds et al., 1983; Martindale et al., 1985; van den Biggelaar, 1996; van den Biggelaar and Guerrier, 1979). As the D quadrant plays a major developmental role in the developing embryo, we wanted to measure the relative sizes of blastomeres in *Maritigrella*, in particular after the second cleavage takes place. Polyclad flatworms, including *M. crozieri*, have been considered equal cleavers on the basis of their indistinguishable relative blastomere sizes at the 2- and 4-cell stages (Lapraz et al., 2013; Martín-Durán and Egger, 2012; Rawlinson, 2010). To test this in *Maritigrella*, we performed a series of precise blastomere volume measurements during the first and second cleavages. We 3D reconstructed 25 fixed embryos between the 2- and 4-cell stages. Additional file 3 (A-E and A’-E’) depicts how the precise volume of given blastomeres can be measured manually using an open source Fiji-plugin (Volumest; http://lepo.it.da.ut.ee/~markkom/volumest/). The measurement data of individual blastomeres can be seen in Additional file 4. For convenience and easier comparison, we labeled vegetal cross-furrow cells in *M. crozieri* as B and D of which the larger cell was always designated as D. Accordingly, the remaining cells were labelled as A and C in consideration of the dextral cleavage type present in *M. crozieri*. One should keep in mind that this assignment may not represent the true quadrants (Rawlinson, 2014) but this process allows us to see at least if there is a consistently larger blastomere and, if so, whether this is an animal or vegetal cross furrow cell.

A small but consistent volume difference of 6% (±1.6%) on average could be discerned between the two blastomeres at the 2-cell stage (n=13) (Figure 5, F and Additional file 4). Two embryos of a transient 3-cell stage show that volumes of the two sister cells also differ (Figure 5, G and Additional file 5) and together have a larger volume than the remaining third blastomere. In the four cell stages, in 9/10 cases, the vegetal cross furrows of the reconstructed embryos were clearly identifiable as schematically drawn in Figure 5, B and depicted in Figure 5, F and F’. Measuring individual blastomeres of 4-cell stage embryos (n = 10) indicates that one of the four cells is larger than the others (Figure 5, F and Additional file 5). This is unlikely to be a random effect as whenever vegetal cross-furrows of 4-cell stage embryos are recognizable, the cell with the largest volume can be identified as one of these. Based on these measurements, *M. crozieri* undergoes asymmetric cell divisions during the first and second cleavages, although they are more pronounced during the two to four-cell transition.

**Figure 5.**
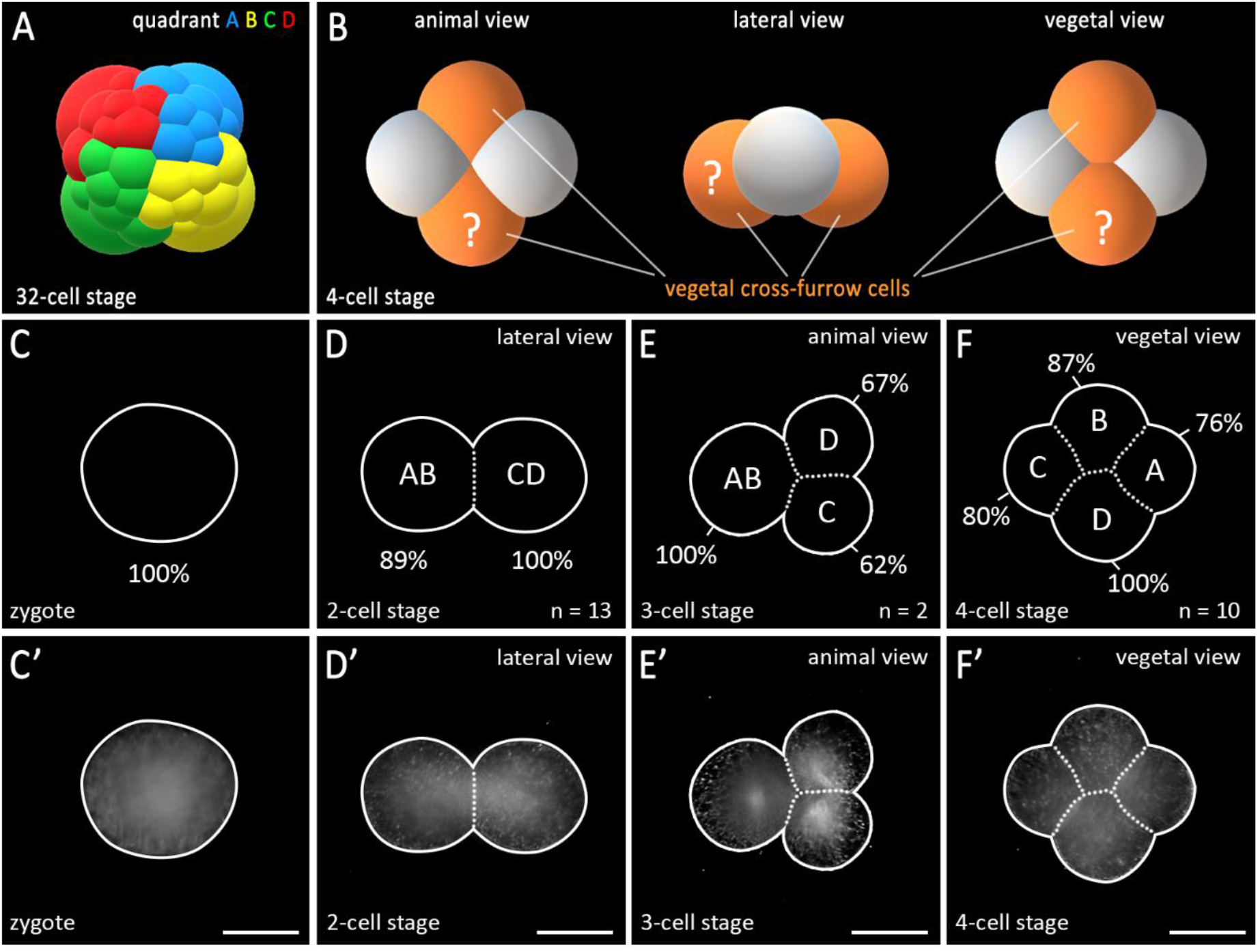
Averaged volume measurements in *M. crozieri* blastomeres of the first and second cleavages. **(A)** A 3D model of a 4-cell stage embryo is depicted showing both vegetal cross-furrow cells that meet at the vegetal pole indicated in orange. **(B)** In unequal cleavers, one of the vegetal cross-furrow cells (depicted here in red) is already specified as the D quadrant at the 4-cell stage. Whether this is true for polyclad flatworms remains unclear, which is indicated here by a question mark. **(C)** A 3D model based on SEM pictures from *M. crozieri* embryos showing the four quadrants (A-D). **(D-H)** Volumes are given as a percentage of the volume of the total embryo, which is 100%. **(F)** At the 2-cell stage the larger cell is assumed to represent blastomere CD and the smaller cell blastomere AB. **(G)** At the 3-cell stage blastomere CD most likely precedes the division of blastomere AB. **(H)** At the 4-cell stage the largest blastomere is always one of the vegetal cross-furrow cells and is interpreted as the D blastomere. **(D’-H’)** All volume measurements come from 5-angle 3D multiview reconstructions and have been orientated with a view from their vegetal side. Only a single plane of the 3D reconstructed stack is shown. Scalebar = 100 μm.

Understanding whether a spiralian embryo is an unequal or equal cleaver is important as it has major implications for determining the mechanism of D quadrant specification. In unequal cleavers (with unequal sized blastomeres at the 4-cell stage), the D quadrant (and therefore the dorsal-ventral axis) can be determined as early as the 4-cell stage: it is assumed that a differential distribution of maternal factors takes place during the first two divisions coinciding with a noticeable inequality of the size of the large D blastomere in comparison to blastomeres A-C (Astrow et al., 1987; Cather and Verdonk, 1979; Clement, 1952; Dorresteijn et al., 1987; Henry, 1986; Henry, 1989; Henry and Martindale, 1987; Render, 1983; Render, 1989).

D-quadrant specification in equal cleavers requires an inductive interaction, usually between one of the equal sized, large vegetal macromeres and the first quartet of micromeres positioned at the animal pole. *So* far, *H. inquilina* is the only polyclad flatworm where earlier blastomere deletion experiments indicated that, in 2-cell and 4-cell stage embryos, asymmetrically distributed morphogenetic determinants could be involved in development (Boyer, 1987) as expected of an unequal cleaver. There is also evidence, however, for the importance of cell-cell interactions between macromeres and micromeres in *H. inquilina* as is typical of equal cleavers (Boyer, 1989).

Our volume measurements indicate that *M. crozieri* does not follow an equal cleavage pattern, although the differences in blastomere size are relatively subtle. At the same time, it is too early to suggest a strictly unequal cleavage mode. It remains possible that there are additional inductive events occurring later in embryogenesis as noted in *H. inquilina*. These could be important for the D quadrant specification and might not be readily observable on a morphological level. In the snail *Illyanassa obsoleta* a mechanism for asymmetric messenger RNA segregation by centrosomal localization during cleavage has been described (Lambert, 2009). It would be very interesting to test for a similar molecular mechanism in polyclad flatworms and to screen for components that play a crucial role in asymmetric cell division machinery as has been recently performed in the spiral cleaving embryo *Platynereis dumerilii* via RNA sequencing (Nakama et al., 2017).

### Micromere 4d in *M. crozieri* shows a cleavage pattern unique to polyclad flatworms

In embryos with spiral cleavage, micromere 4d typically divides into a left and a right daughter cell by a meridional division. It is at this point that the bilateral symmetry of the embryo first emerges at a cellular level. To determine the symmetry breaking event during *M. crozieri* development we followed the division pattern of micromere 4d using our live-imaging data. We observe that the 4d blastomere in *M. crozieri* does not divide meridionally into a left and right daughter cell, but first divides along the animal-vegetal axis into a smaller, animally positioned cell, which we designate as 4d^2^ and a larger, vegetally positioned cell, we designate 4d^1^ (Figure 6, A-B and F-H; Additional file 6). We thereby follow closely the nomenclature used by Surface (1907) and it should be noted that in this specific case (the animal-vegetal division of an ento- and mesoblast and not the ectoblast) the smaller exponent was intentionally reserved for the more vegetally positioned “parent” cell. Only following this additional division of micromere 4d, is definitive bilateral symmetry established by the meridional (left-right) division of both sister cells, 4d^1^ and 4d^2^ (Figure 6, C-E and H-J). The meridional divisions of 4d^1^ and 4d^2^ appear equal and this equality is easily observed in 4d^2^ due to its larger size and exposed external position. Both descendants of 4d^1^ and 4d^2^ (4d^11^ and 4d^12^ and 4d^21^ and 4d^22^) then undergo another round of roughly meridional cleavages. This is similar to Surface’s descriptions in *H. inquilina* (Surface, 1907).

**Figure 6.**
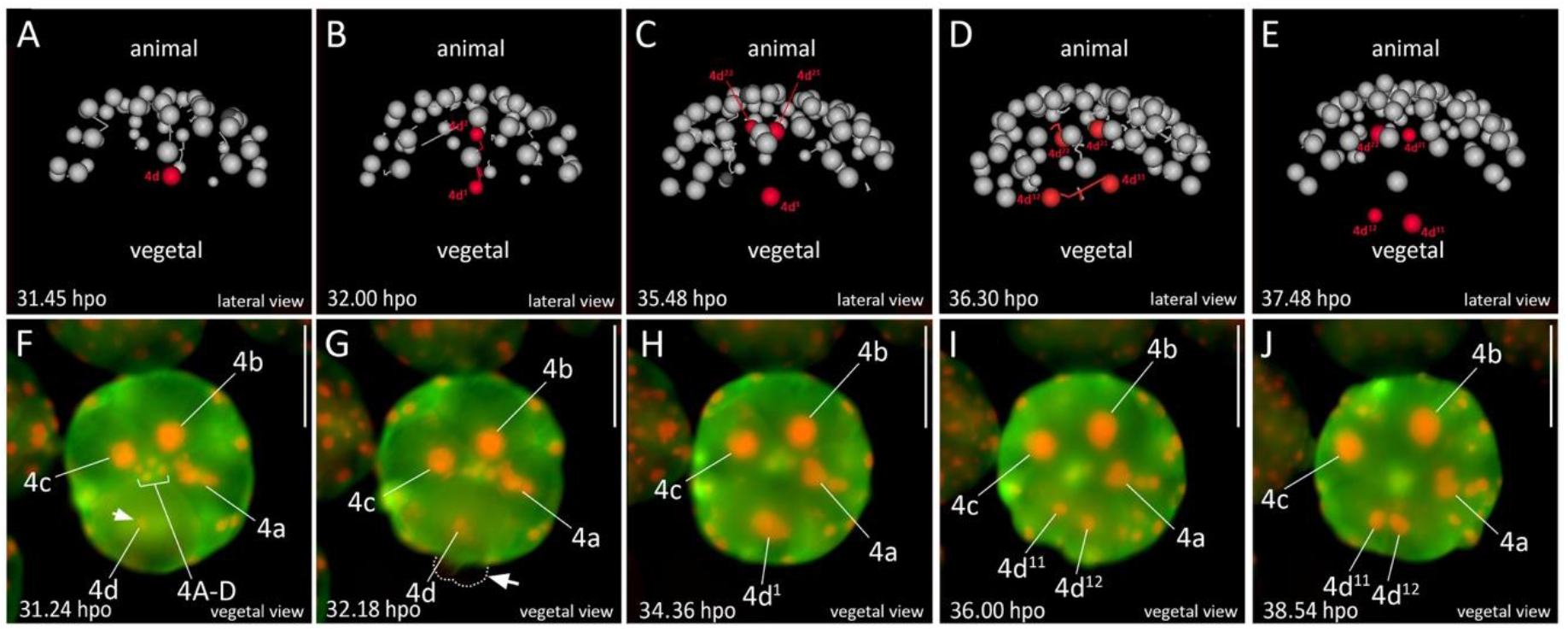
Animal view of the cleavages of micromere 4d in *M. crozieri*. **(A-E)** The cleavage pattern of micromere 4d (marked in red) is visualized using a 3D viewer (Fiji), showing in grey the position of all remaining nuclei except 4A-D and 4a-4c. **(A)** Micromere 4d before its division. **(B)** Micromere 4d divides along the animal-vegetal pole and daughter cell 4d^2^ is budded into the interior of the embryo and in close proximity to micromeres of the animal pole. **(C-E)** Both daughter cells of micromere 4d divide again, but this time both cells cleave meridionally **(F)** Micromere 4d undergoes mitosis revealing the D quadrant. **(G)** The asymmetric division of micromere 4d along the animal-vegetal pole is barely visible but causes blebbing (arrow pointing at dashed line). **(H)** After the division, daughter cell 4d^1^ remains large and is more vegetally positioned and therefore readily visible. 4d^2^ is budded into the interior of the embryo, more animally positioned and cannot be seen anymore without optical sectioning. **(I-J)** Bilateral symmetry is clearly visible after the division of 4d^1^. Oocytes were microinjected with nuclear marker H2A-mCherry (red) and microtubule marker EMTB-3xGFP (green) and the embryo used for 4d microscopy with OpenSPIM (A-E) or under a Zeiss Axio Zoom.V16 Stereo Microscope (F-J); hpo = hours post oviposition. Scalebar in images captured with the Axio Zoom = 100 μm.

Surface (1907) and later van den Biggelaar (1996) both already noted that in polyclads the cleavage of 4d differs from the canonical pattern of an immediate, equal and meridional division into left and right descendants. According to van den Biggelaar, in the polyclads *Hoploplana inquilina* and *Prostheceraeus giesbrechtii*, this meridional division is delayed by one cell cycle as 4d first undergoes the approximately animal-vegetal division into 4d^1^ and 4d^2^. This is followed by meridional cleavages of both daughter cells 4d^1^ and 4d^2^. These observations exactly match what we observe in *M. crozieri*. In other more recent descriptions of polyclad flatworms (Hartenstein and Ehlers, 2000; Malakhov and Trubitsina, 1998; Rawlinson, 2010; Teshirogi et al., 1981; Younossi-Hartenstein and Hartenstein, 2000), this the animal-vegetal division of 4d is not mentioned suggesting either that some polyclad flatworms lack it or, more likely, that the division is difficult to observe without continuous recording. Our observations in the *M. crozieri* together with description of *H. inquilina* by Surface and *P. giesbrechtii* by Van den Biggelaar strongly suggests that this cleavage pattern of micromere 4d is in fact unique amongst spiralians but common across polyclads.

### Post-meiotic protrusions of the cell membrane (blebbing) accompany early development in *M. crozieri*

In several animal phyla, oocytes undergo cytoplasmic changes that are capable of temporarily deforming the shape of the egg and which have been suggested as a sign of the oocyte segregating cell content (Wall, 1990). Such events have been commonly observed during fertilization and meiosis (Henry et al., 2006; Lehmann and Hadorn, 1946; Li and Albertini, 2013; Meshcheryakov, 1991). In polyclads this has been demonstrated many times previously and is referred to as cell blebbing (Anderson, 1977; Gammoudi et al., 2011; Goette, 1881; Hallez, 1879; Kato, 1940; Malakhov and Trubitsina, 1998; Rawlinson et al., 2008; Selenka, 1881; Surface, 1907; Teshirogi et al., 1981; Younossi-Hartenstein and Hartenstein, 2000). It has occasionally been noted that cell-blebbing is not restricted to egg maturation and the extrusion of the polar bodies but can reappear frequently during early cleavages ((Gammoudi et al., 2012; Malakhov and Trubitsina, 1998; Teshirogi et al., 1981).

In *M. crozieri*, our observations show that blebbing during egg maturation follows first a depression of the oocyte at the animal pole (Figure 7, A) followed by protrusions all over the cell membrane (Figure 7, C and insets). These events are almost identical to drawings of egg maturation and oocyte blebbing based on different Japanese polyclad species by Kato (1940).

**Figure 7.**
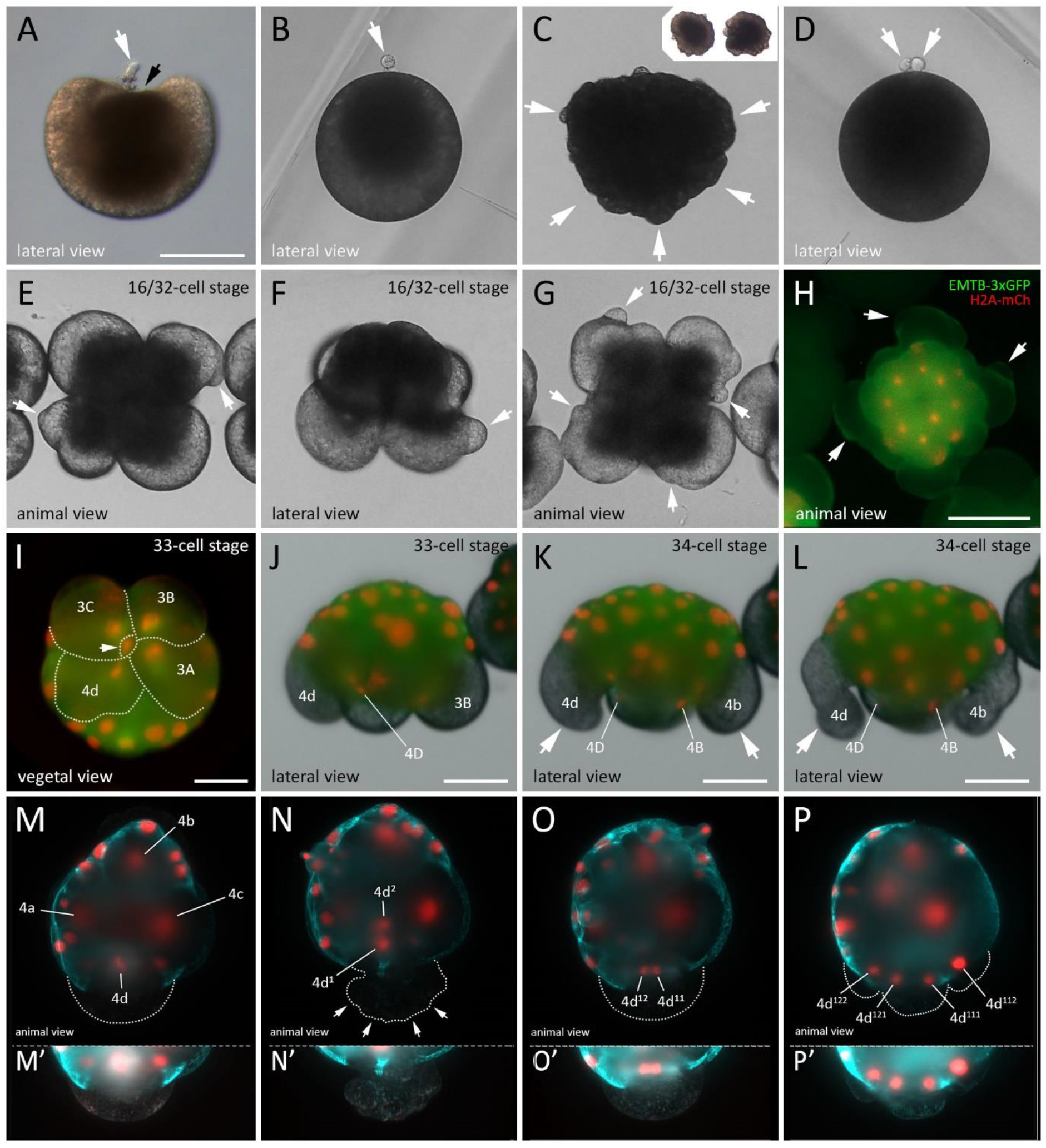
Blebbing events during meiosis and spiral cleavage in the polyclad flatworm M. crozieri **(A-D)** Blebbing during egg maturation in *M. crozieri* oocytes. **(A)** Extrusion of first polar body (white arrow) and depression of the oocyte at the animal pole (black arrowhead). **(B)** Oocyte with one polar body and darkish pigment accumulated at the animal pole **(C)** Cell blebbing is recognisable by the formation of amoeboid/pseudopodia-like irregularities all over the cell membrane. **(D)** Egg cell with two polar bodies and darkish pigment accumulated at the animal pole. **(E-L)** Blebbing during the third and fourth quartet formation **(E-H)** Protrusions in the form of extracellular vesicle-like structures appear prior to third quartet formation (16-32-cell stage) among all four macromeres. **(I-J)** Vegetal **(I)** and lateral view **(J)** of the division of macromere 3D into tiny macromere 4D (white arrowhead). **(K-L)** Blebbing is accompanied by severe deformations of large micromeres 4b and 4d. **(M-P)** Animal view of the cleavages of micromere 4d in *M. crozieri*. **(M)** Chromosome condensations are only visible in 4d. **(N)** Division of 4d is visible along the animal-vegetal axis of the embryo. White arrowheads show cytoplasmic perturbations during the cleavage of micromere 4d. **(O)** Meridional division of 4d^1^ takes place. **(P)** The next division of the daughter cells of 4d^1^ is depicted. **(M’-P’)** The 4d-cell and its progenies have been depicted separately below at increased exposure levels. Embryos with fluorescent signal were microinjected as oocytes with a microtubule marker (EMTB-3xGFP) and a histone nuclear marker (H2A-mCh). Live imaging was performed under a under a Leica DMI3000 B inverted scope (A-G), a Zeiss Axio Zoom.V16 Stereo Microscope (H-L) and an OpenSPIM (M-P). Scalebar is 100 μm in A and H, 50 μm in I-L, 100 μm in A and E and 50 μm in M-P.

Blebbing in *Maritigrella* continues after meiosis, specifically during the asymmetric cleavages of macromeres (Figure 7). The formation of the third and fourth quartet micromeres is clearly accompanied by strong blebbing events in the macromeres distinct from what is seen in meiotic cell blebbing (Figure 7, E-L). In the case of the third quartet formation we observe that, prior to the cleavage of macromeres 2A-2D, blebbing becomes visible on their cell surfaces in form of small, vesicle-like protrusions (Figure 7, E-H) (n = 17/18). The role of these vesicles is not clear, but we can observe that mitotic cytoskeletal activity during anaphase correlates with the observed protrusions (Additional file 7 and Additional file 8). In contrast, during the formation of the fourth quartet (3A-3D), cytoplasmic perturbations create waves of contractile activity with smaller blebs that appear more frequently. In this case, the macromeres can sometimes attain an elongated shape (Figure 7, I-L) (n=18/18) at the onset of the formation of micromeres 4a-4d. More detailed time-lapse sequences of these peculiar cytoplasmic perturbations are shown in Additional file 9. Finally, the asymmetric division of micromere 4d in *M. crozieri* is also accompanied by distinctive cytoplasmic perturbations of the membrane (Figure 7, M-P and M’-P’); n = 16/16). The perturbations of micromere 4d mark the end of a series of cell shape changes visible throughout early development. In Figure 8 we summarise the events previously described in polyclad flatworms, together with our own observations of the early development of *M. crozieri*.

**Figure 8.**
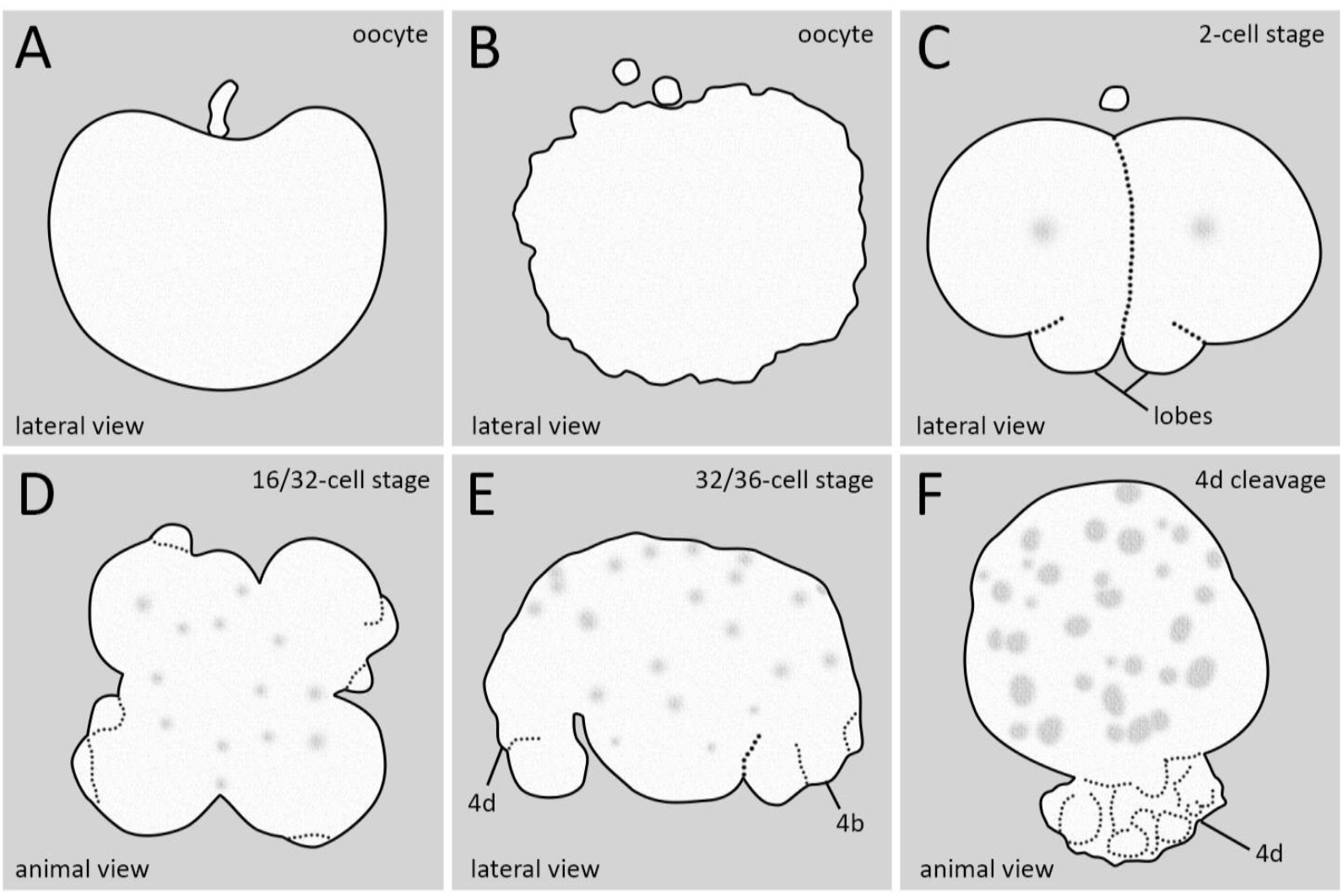
Summary of cytoplasmic perturbations described in different polyclad flatworm species. **(A)** Depression of the animal pole during the formation of the first polar body described by Kato (1940) for some Japanese polyclad species and for *Maritigrella crozieri* (this study). **(B)** Cell blebbing in oocytes as described for most polyclads during the first and second meiotic divisions (see Gammoudi et al. 2012). **(C)** Vegetal lobe like structures found in *Pseudostylochus intermedius* (Teshirogi & Sachiko, 1981) and *Pseudoceros japonicus* (Malakhov and Trubitsina, 1998). Drawing taken from *P. intermedius* **(D)** Cytoplasmic perturbations seen in *Pseudostylochus intermedius* (8-to 16-cell stage) (Teshirogi & Sachiko, 1981) and *Maritigrella crozieri* (16-to 32-cell stage, this study). **(E)** Waves of contractile activity in all four macromeres of *Maritigrella crozieri* (this study) whereby macromeres attain an elongated shape. **(F)** Similar cytoplasmic perturbations seen during the highly asymmetric cleavage of micromere 4d found in *Maritigrella crozieri* (this study).

At present it is unclear what role (if any) post-meiotic blebbing plays during early cleavage. It is interesting to see that the perturbations observed in *Maritigrella* during divisions of macromeres 2A-2D (extracellular vesicle-like structures; see Figure 7, E-H) look identical to a highly similar blebbing event in another polyclad species, the acotylean *Pseudostylochus intermedius* (Teshirogi et al., 1981), although in the latter species this phenomenon is described to take place one division round earlier (8-to 16-cell stage). Blebbing during the divisions of macromeres 3A-3D and the division of micromere 4d are both described by us for the first time during polyclad embryogenesis.

One observation suggesting blebbing has an important function in polyclad embryogenesis is that when embryos are mounted in high concentrations of agarose (>0.6%) we observed severely abnormal development (n=5/5). We speculate that these defects may be caused by blebbing being hampered by the stiff agarose. Common to all of the blebbing events is that they occur in cells which contain most of the yolk and which undergo asymmetric divisions. Additionally, we show here that they coincide with increased cytoskeletal activity (mitosis). One simple explanation for their occurrence may be that they are the visible manifestation of actomyosin contractions of the cortex during strong cytoskeletal movements involved in asymmetric cleavage in yolk-rich blastomeres.

## Conclusions

In this study we have used live-imaging recordings and 3D reconstructions to extend observations of early development in a cotylean polyclad flatworm, *Maritigrella crozieri*. 3D reconstructions and continuous 4D recordings allow us to see developmental events in more detail than previously possible. We have been able to look at connections between nuclear movements and cell divisions and link them with cellular dynamics such as cell blebbing (protrusions of the membrane), and pinpoint important developmental events like symmetry breaking. Our observations allow us to confirm and extend previous developmental observations of early embryogenesis in polyclads, made using fixed specimens, describing the spiral cleavage pattern and the formation of the four quartets. There seems to be little variation within both polyclad suborders, the cotyleans and the acotyleans.

One important observation in *M. crozieri* is that this so-called equal cleaving polyclad should probably not be classified as such. Our measurements of individual blastomeres at the 4-cell stage show that the second cleavage is a product of unequal divisions of which one vegetal cross-furrow blastomere retains the largest volume. Similar observations of unequal cleavage may be a broader pattern within polyclads, including species so far regarded as “equal” cleavers, but requires precise measurements to be carried out in different species. In *M. crozieri* the question remains as to whether the observed size differences at the 4-cell stage truly reflect an unequal cleavage mechanism, meaning that the D quadrant is already specified by maternal determinants at this early stage. Clearly, we need to know more about the molecular basis of putative maternal determinants and the mechanisms by which they could be sequestered but an early specification of the D quadrant via cytoplasmic localization seems to be supported by previous studies on *Hoploplana inquilina* (Boyer, 1987; Boyer, 1989).

Most importantly, we found that the animal-vegetal division of micromere 4d is present in both polyclad suborders, and we suggest this is a conserved pattern across all polyclad flatworms. It would be highly interesting to reinvestigate this cleavage pattern within the Prorhynchida, where the spiral cleavage pattern with quartet formation has also been partly retained but current developmental data are insufficient to conclude whether it follows the pattern as suggested for polyclads in this study.

We consider that the exact fate of both daughter cells of micromere 4d must be investigated more thoroughly before we can conclude whether micromere 4d^2^ (animally positioned relative to 4d^1^) indeed represents the mesentoblast or not. Currently, even the fate of 4d^1^, despite its large size and the fact that it is readily visible at the onset of gastrulation, remains unclear, as model lineage tracing of this specific blastomere has not been yet performed. This could be done by DiI injections or perhaps via fluorescently tagged and photoconvertible molecules. It would also be interesting to study further the apparent interaction of one of the daughter cells 4d^1^ with small macromeres (4A-D), observed during our live-imaging recordings in *M. crozieri*, as this is the first time that cell-cell interactions may have been directly identified in a polyclad flatworm and that a potentially significant developmental role is assigned to the small macromers.

We show here new evidence that, in *M. crozieri*, blebbing is present not only in oocytes during meiosis, but also in macromeres during quartet formation and in micromere 4d during its first cleavage along the animal-vegetal axis (Figure 6, Figure 7 and Additional file 6). We propose that it is likely that these are a manifestation of the mechanical forces created by cytoskeletal dynamics during early cleavages, which may be more or less obvious depending on the polyclad species and the amount of yolk within the blastomere. Alternatively, these movements could fulfill other purposes such as correctly sequestering factors that could play a role in development, but this remains to be seen in future studies.

Taken together the most crucial events during polyclad spiral cleavage take place as follows: Firstly a D quadrant might be established as early as the 4-cell stage by cytoplasmic localizations (Boyer, 1989). The atypical formation of the fourth quartet then gives rise to micromere 4d, which arguably behaves similarly to macromere 3D in molluscan and annelid embryos (van den Biggelaar, 1996). Unusually, micromere 4d undergoes an animal-vegetal division, which buds micromere 4d^2^ into the interior of the embryo and in proximity to the animal cap as shown by Surface (1907) in *H. inquilina* and *M. crozieri* (this study). In our opinion the position 4d^2^ assumes during this event allows it the possibility to interact with micromeres of the first quartet. Such animal-vegetal inductive interactions are typically observed in equally cleaving spiralians (Lyons and Henry, 2014) and could also play a crucial role in polyclads in terms of specifying the D quadrant. Ultimately, 4d^2^ may be considered the mesentoblast (Martín-Durán and Egger, 2012), but this still remains to be determined more carefully.

As shown in this work, polyclad flatworms appear to combine conserved features of spiral cleavage but also show obvious modifications of their cleavage program. This makes them a highly interesting taxa for evolutionary comparisons among flatworms within and outside the polyclad order but also across lophotrochozoan phyla. Live imaging recordings such as SPIM can certainly contribute also in future studies to extend our current understanding of polyclad development and other marine invertebrates.

## Methods

### Animal culture

Adult specimens of M. crozieri were collected in coastal mangrove areas in the Lower Florida Keys, USA in January 2014, November 2014 September 2015 and January 2016 near Mote Marine Laboratory (24.661621, −81.454496). Animals were found on the ascidian *Ecteinascidia turbinata* as previously described (Lapraz et al., 2013). Eggs without egg-shells (to produce ‘naked’ embryos) were obtained from adults by poking with a needle (BD Microlance 3) and raised in Petri dishes coated with 2% agarose (diluted in filtered artificial seawater) or gelatin coated Petri dishes at room temperature in penicillin-streptomycin (100 μg/ml penicillin; 200 μg/ml streptomycin) treated Millipore filtered artificial seawater (35-36 ‰).

### *In vitro* synthesis of mRNA

We synthesised mRNAs for microinjections with Ambion’s SP6 mMESSAGE mMACHINE kit. The capped mRNAs produced were diluted in nuclease-free water and used for microinjections in order to detect fluorescence signal in early *M. crozieri* embryos. Nuclei were marked and followed using histone H2A-mCherry (H2A-mCh) and GFP-Histone (H2B-GFP). The plasmids carrying the nuclear marker pCS2-H2B-GFP (GFP-Histone) and pDestTol2pA2-H2A-mCherry (Kwan et al., 2007) were transformed, purified and concentrated as described before and then linearized with the restriction enzymes NotI and BglII respectively. To follow live microtubules, we used a GFP fusion of the microtubule binding domain of ensconsin (EMTB-3XGFP). These clones were the gift of the Bement Lab (University of Wisconsin) (Burkel et al., 2007; Miller and Bement, 2009) and were commercially ordered from http://addgene.org (EMTB-3XGFP: https://www.addgene.org/26741/).

### Microinjections

Fine-tipped microinjection needles were pulled on a Sutter P-97 micropipette puller (parameters: P=300; H=560; Pu=140; V=80; T=200.) and microinjections of synthesized mRNA (~300-400 ng/μl per mRNA in nuclease-free water) were carried out under a Leica DMI3000 B inverted scope with a Leica micromanipulator and a Picospitzer^®^ III at room temperature.

### 4D microscopy of live embryos using OpenSPIM

Embryos showing fluorescent signal were selected under an Axioimager M1 Epifluorescence and Brightfield Microscope (Zeiss). Live embryos were briefly incubated in 40 °C preheated and liquid low melting agarose (0.1%) and immediately sucked into fluorinated ethylene propylene (FEP) tubes (Bola S1815-04), which were mounted in the OpenSPIM acquisition chamber which was filled with filtered artificial seawater and antibiotics via a 1 ml BD Plastikpak (REF 300013) syringe. The use of FEP tubes has been previously described (Kaufmann et al., 2012) and allows the specimen to remain inside the tube during image acquisition without causing any blurring to the acquired images, as would be the case with other mounting materials such as glass capillaries. Using FEP tubes enables us to mount specimens in lower percentage agarose (0.1-0.2%), thus lessening the perturbation of embryo growth and development. The interval between images depends on the user’s intentions. Long-term imaging single timepoints can consist of 40-70 optical slices and were captured every 1-3 minutes. The OpenSPIM was assembled according to our previous description (Girstmair et al., 2016) and operated using MicroManager (version 1.4.19; November 7, 2014 release; https://www.micro-manager.org/).

### 4D microscopy of live embryos under an Axio Zoom.V16 (Zeiss)

Several embryos in which fluorescent signal could be detected were centered within a 90 mm petri dish containing penicillin-streptomycin (100 μg/ml penicillin; 200 μg/ml streptomycin) treated Millipore filtered artificial seawater (35-36 ‰) for simultaneous live imaging. To avoid evaporation and make fluorescent imaging possible a tiny hole was made in the middle of the lid and artificial seawater containing fresh antibiotics carefully exchanged from the side when evaporation became apparent. Brightfield, green and red fluorescence was acquired every 5-7 min.

### Fixation and imaging of embryos used for scanning electron microscopy (SEM)

Batches of embryos were raised until development reached the desired stage (1-cell, 2-cell, 4-cell, 8-cell, 16-cell, 32-cell, 64-cell and intermediate phases). Fixation was done at 4°C for 1 hour in 2.5% glutaraldehyde, buffered with phosphate buffered saline (PBS; 0.05 M PB/0.3 M NaCl, pH 7.2) and post-fixed at 4°C for 20 min in 1% Osmium tetroxide buffered with PBS. Fixed specimens were dehydrated in an ethanol series, dried via critical point drying, and subsequently sputtered coated with carbon or gold/palladium in a Gatan 681 High Resolution Ion Beam Coater and examined with a Jeol 7401 high resolution Field Emission Scanning Electron Microscope (SEM).

### Fixation and staining of embryos for 3D reconstruction

Embryos were extracted from gravid adults at the Keys Marine Laboratory (Florida) by poking and allowed to cleave until the desired stage was reached. Embryos were then fixed for 60 min in 4 % formaldehyde (from 16 % paraformaldehyde: 43368 EM Grade, AlfaAesar) in PBST (0.1 M phosphate buffer saline containing 0.1% Tween 20) at room temperature, followed by a 5x washing step in PBST and stored at 4 °C in PBST containing small concentrations of sodium azide.

In order to image specimens from 5 angles, which is necessary to perform volume measurements of early blastomeres, sodium azide with 0.1 M PBS containing 0.1% Triton X-100 in (PBSTx) was washed off fixed embryos by four washing steps and stained with 1:300 Rhodamine Phalloidin (ThermoFisher Scientific R415) for 2-3 h at room temperature or overnight at 4°C. Following several washes of PBST or PBSTx 0.1 μM of the nuclear stain SytoxGreen (Invitrogen), which is difficult to detect at these early stages, was added for 30 min and the embryos then rinsed with PBST for another hour.

### Image processing

Post-processing of acquired data was performed with the latest version of the freely available imaging software Fiji (Schindelin et al., 2012) and digital images were assembled in Adobe Photoshop CC 2017.

### Ethics approval and consent to participate

Not applicable.

### Consent for publication

Not applicable.

### Availability of data and materials

The datasets during and/or analysed during the current study available from the corresponding author on reasonable request.

### Funding

JG was funded by the Marie Curie ITN ‘NEPTUNE’ grant (no. 317172), under the FP7 of the European Commission. M.J.T. was supported by the European Research Council (ERC-2012-AdG 322790), the Biotechnology and Biological Sciences Research Council grant (BB/H006966/1 and a Royal Society Wolfson Research Merit Award.

## Author contributions

MJT and JG designed the experiments. JG performed all the experiments and prepared the figures. JG and MJT analysed the data and wrote the manuscript. The authors read and approved the final manuscript.

## Acknowledgements

We thank Anne Zakrzewski for her help with scanning electron microscopy pictures, Bernhard Egger and Kate Rawlinson for their continuous support and helpful discussions, Fraser Simpson for his immense effort during the Florida collection trips, the Mote Marine Laboratory (https://mote.org/) and Keys Marine Laboratory (http://www.keysmarinelab.org/) for their support during our stay in the Florida Keys.

## List of abbreviations

(FEP): fluorinated ethylene propylene
(MTOC): microtubule organizing centre
(SEM): scanning electron microscopy
(SPIM): selective plane illumination microscopy
(SD): spiral deformations
(PBS): phosphate buffered saline

## ADDITIONAL FILES

Additional file 1 – 50 min OpenSPIM movie of the third cleavage in an embryo of *M. crozieri* with labeld nuclei (H2B:GFP) showing spiral deformations (SD) and dexiotropic cleavage.

Additional file 2 – Putative cell-cell interactions captured with an OpenSPIM of an embryo undergoing epiboly. It can be observed how nuclei of the small macromeres (4A-4D) get in very close proximity with nuclei of close descendants of micromere 4d^2^ for a short period of time and then goes away.

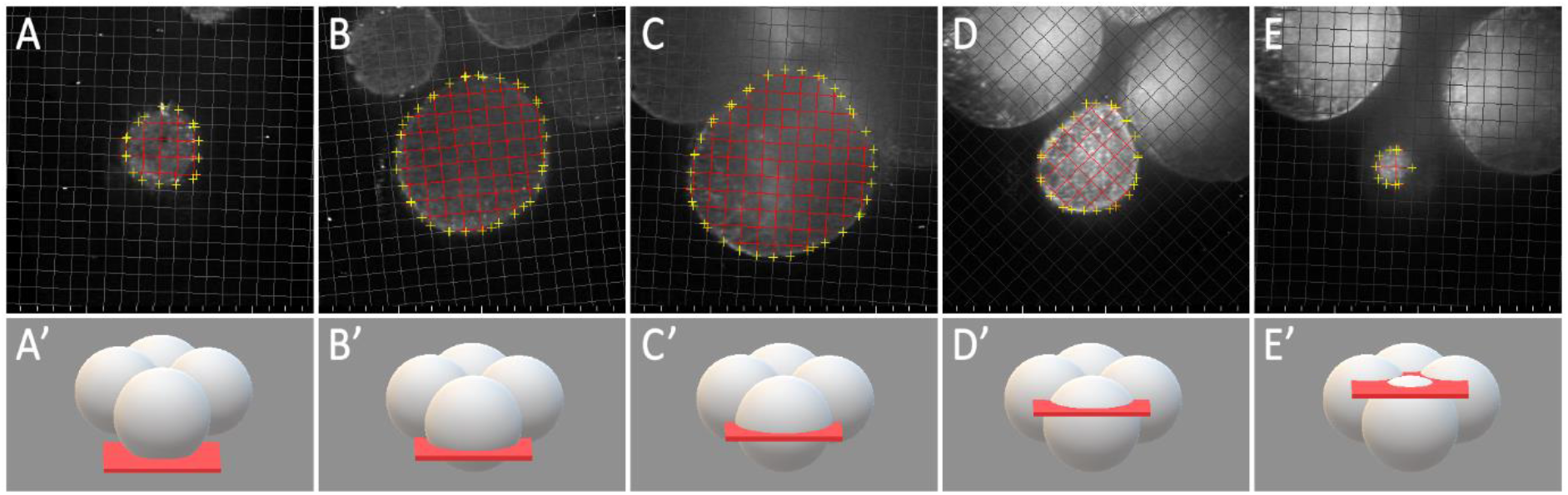

Additional file 3 – An example of volume measurements performed on a 4-cell stage polyclad flatworm embryo, showing only 5 representative slices within a Z-stack (the original file contains hundreds of slices after image processing is completed).

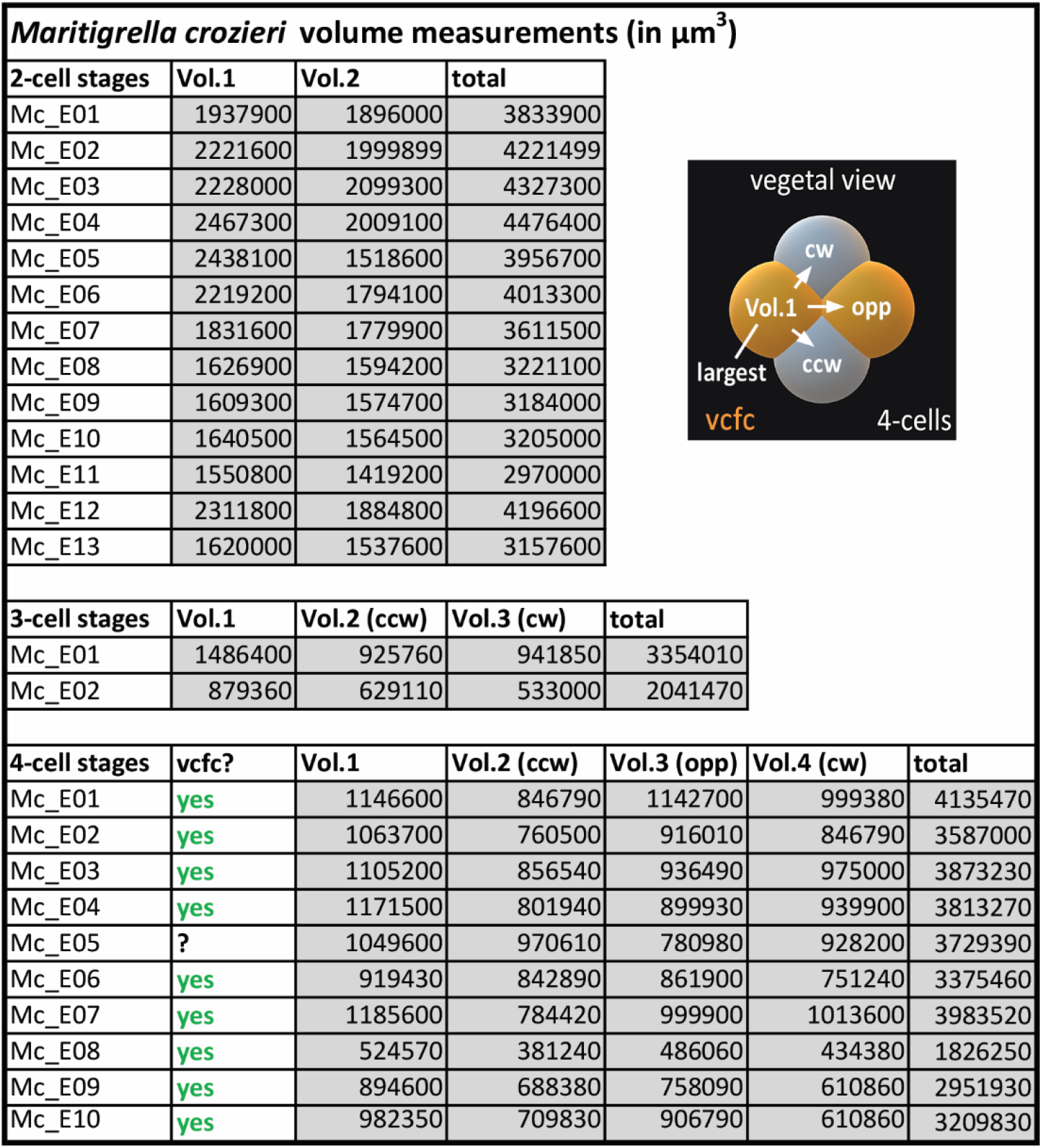

Additional file 4 – Table of blastomere volume measurements in 2-, 3- and 4-cell stages. Vol.1 indicates the largest blastomere. In 2-cell stages Vol.2 accounts for its sister cell. In 4-cell stages Vol.2 corresponds to cells positioned clockwise (cw) of it, Vol. 3 to the cell opposite of it (opp) and Vol.4 counter clockwise (ccw) of it (see schematic embryo inset). The vegetal cross-furrow-cells (vcfc) are shown in orange.

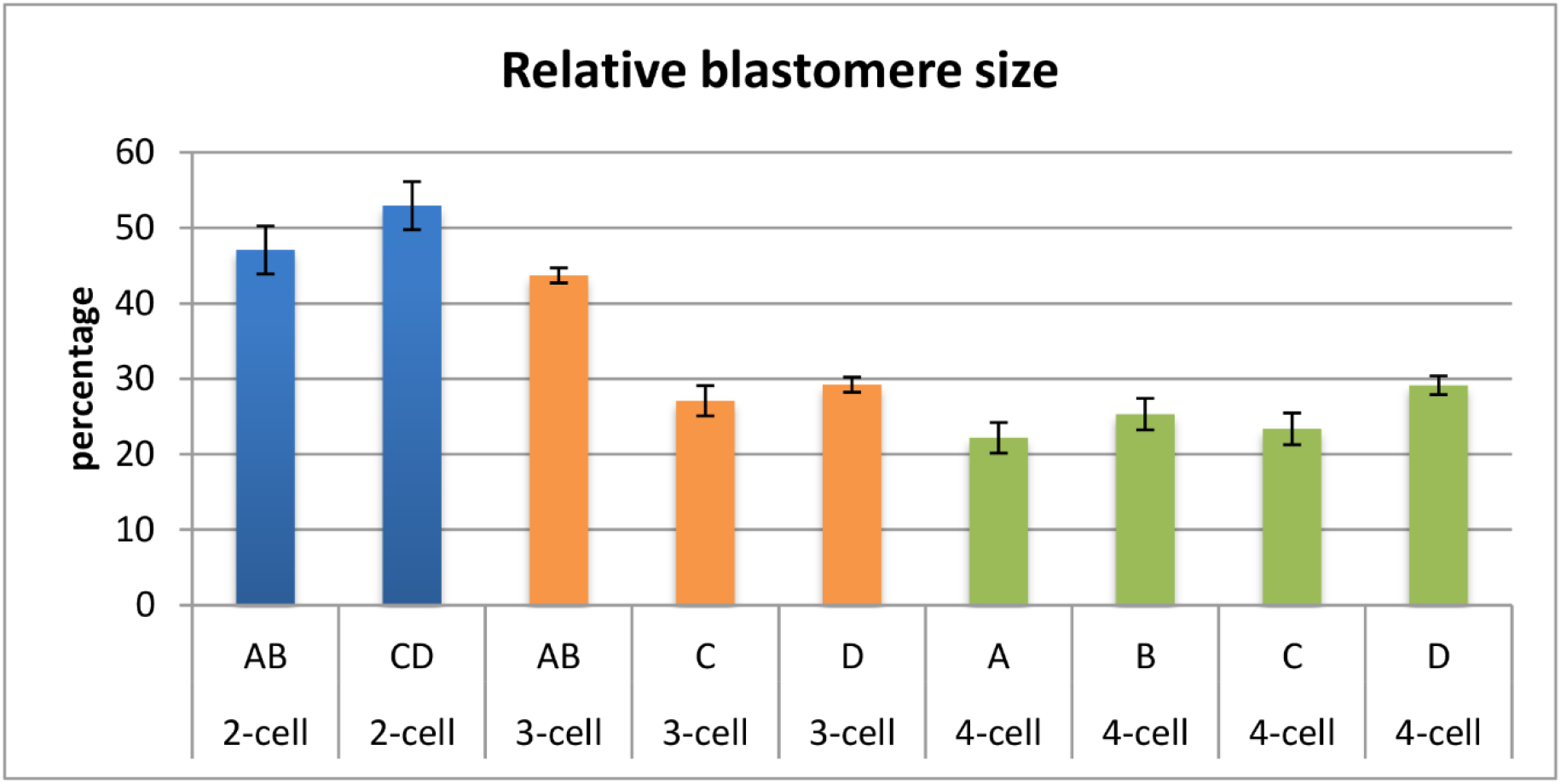

Additional file 5 – An average of the volume measurements of 3D reconstructed blastomeres in *M. crozieri* embryos of the 2-cell, 3-cell and 4-cell stages. The data are based on measurements of individual blastomeres. To provide the data as percentages makes sense as each individual embryo can vary in size. The 2-cell stages are indicated as blue columns (n=13), 3-cell stages as orange (n=2) and 4-cell stages as green columns (n=13). Volumes are given as a percentage of the total volume of the embryo which is 100%. Standard deviations are indicated for smaller blastomeres only. In two-cell stages a 6% difference was noted between the two cells on average. The larger blastomere has been designated as CD. In 3-cell stages the two sister blastomeres (C and D) have a larger volume than the remaining sister cell and have been designated as C and D according to a slight volume difference. In 4-cell stages the largest blastomere is one of the vegetal cross-furrow cells and has been indicated as D. It is 5.8% larger compared to its sister cell indicated as C. Of the two, remaining sister blastomeres, the size difference is only 3.3% with the larger one indicated as blastomere B. Error bars indicate standard error of the mean.

Additional file 6 – The initial division pattern of micromere 4d using live-imaging data from an Axio Zoom.V16 (Zeiss). The 4d blastomere does not divide laterally but first divides along the animal-vegetal axis into a smaller, animally positioned cell, which we designate as 4d^1^ and a larger, vegetally positioned cell, we designate 4d^2^

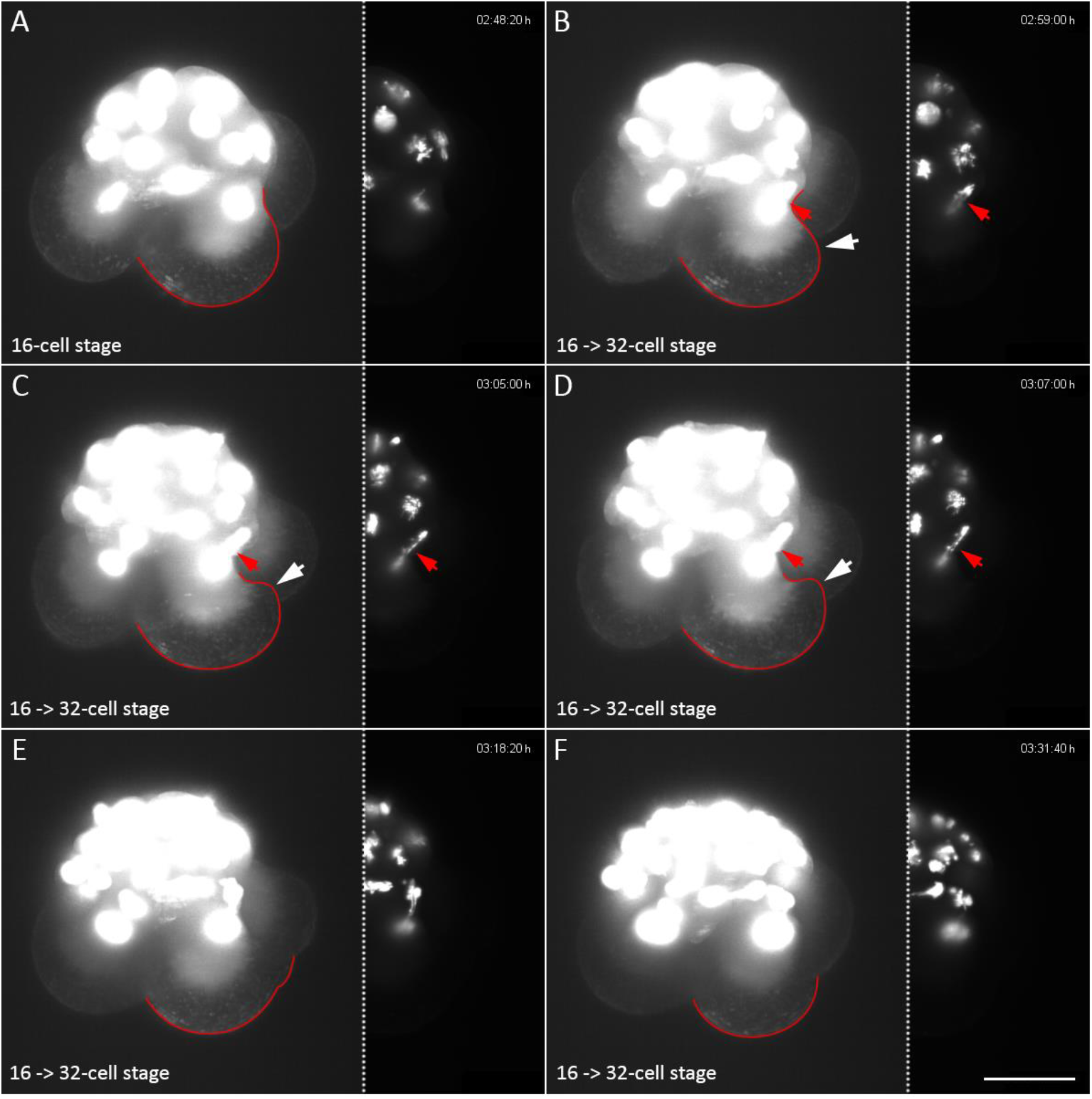

Additional file 7 – Cytoplasmic perturbations imaged with the OpenSPIM in one of the second quartet macromeres shown during mitosis. The whole embryo is shown over-exposed to better visualize the membranous outlines of the macromeres. To the right of each embryo, the nuclei are depicted with normal exposure. Red arrows point to the same nucleus of the embryo. A red line highlights the outline of the corresponding macromere. The shape deformations caused by the cytoplasmic perturbations of the macromere correlate precisely with the mitotic anaphase and reach a maximum in panel D. Scalebar = 50 μm.

Additional file 8 – Movie of and embryo forming the third quartet. Prior to the cleavage of macromeres 2A-2D, blebbing becomes visible on their cell surfaces in form of small, vesicle-like protrusions. The movie shows that mitotic cytoskeletal activity during anaphase correlates with the observed protrusions.

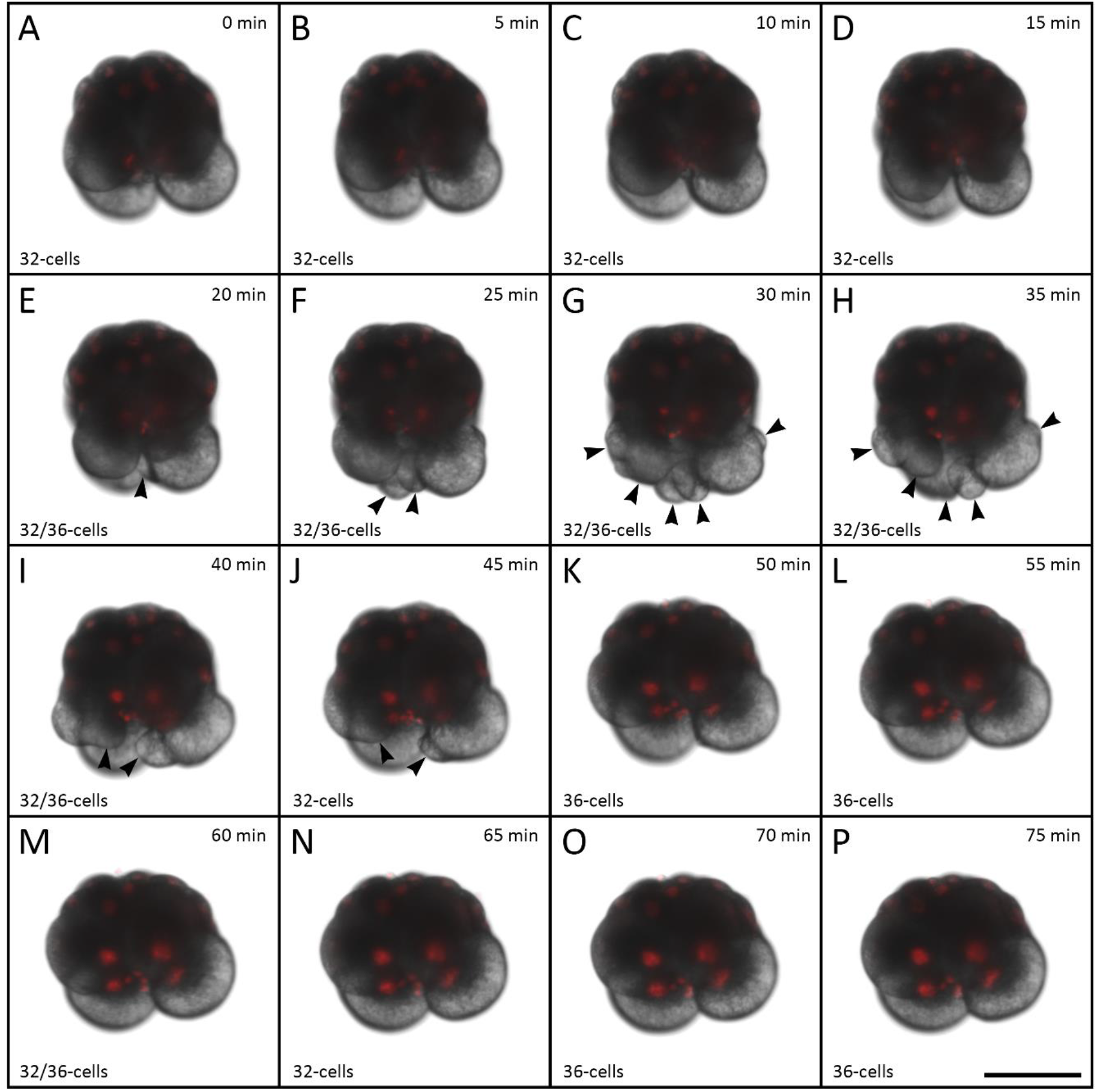

Additional file 9 – **(A-P)** Time-lapse recording showing the formation of the small macromeres (4Q) and large micromeres (4q) of a single *M. crozieri* embryo in 5 min intervals showing striking cytoplasmic perturbation activity at the vegetal pole of the embryo (indicated by black arrows). **(F-K)** 25 min of cytoplasmic perturbations are clearly visible in macromeres 3A-3D. Live imaging was performed under a Zeiss Axio Zoom.V16 Stereo Microscope. Scale bar is 100 μm.

